# Repurposing Ribo-Seq to provide insights into structured RNAs

**DOI:** 10.1101/2020.05.18.103374

**Authors:** Brayon J. Fremin, Ami S. Bhatt

## Abstract

Ribosome profiling (Ribo-Seq) is a powerful method to study translation in bacteria. However, this method can enrich RNAs that are not bound by ribosomes, but rather, are protected from degradation in another way. For example, *Escherichia coli* Ribo-Seq libraries also capture reads from most non-coding RNAs (ncRNAs). These fragments of ncRNAs pass all size selection steps of the Ribo-Seq protocol and survive hours of MNase treatment, presumably without protection from the ribosome or other macromolecules or proteins. Since bacterial ribosome profiling does not directly isolate ribosomes, but instead uses broad size range cutoffs to fractionate actively translated RNAs, it is understandable that some ncRNAs are retained after size selection. However, how these ‘contaminants’ survive MNase treatment is unclear. Through analyzing metaRibo-Seq reads across *ssrS*, a well established structured RNA in *E. coli*, and structured direct repeats from Clustered Regularly Interspaced Short Palindromic Repeats (CRISPR) arrays in *Ruminococcus lactaris*, we observed that these RNAs are protected from MNase treatment by virtue of their secondary structure. Therefore, large volumes of data previously discarded as contaminants in bacterial Ribo-Seq experiments can, in fact, be used to gain information regarding the *in vivo* secondary structure of ncRNAs, providing unique insight into their native functional structures.

**Importance:** We observe that ‘contaminant’ signals in bacterial Ribo-Seq experiments that are often disregarded and discarded, in fact, strongly overlap with structured regions of ncRNAs. Structured ncRNAs are pivotal mediators of bioregulation in bacteria and their functions are often reliant on their specific structures. We present an approach to access important RNA structural information through merely repurposing ‘contaminant’ signals in bacterial Ribo-Seq experiments. This powerful approach enables us to partially resolve RNA structures, identify novel structured RNAs, and elucidate RNA structure-function relationships in bacteria at a large-scale and *in vivo*.

## Observation

Ribosome profiling (Ribo-Seq) in bacteria is a method that enriches for ribosome-protected RNAs and therefore, enables the study of active translation events^1,2^. However, this enrichment is not highly selective for ribosomes as relevant protocols do not specifically isolate ribosomes, but rather select for them within an expected size range. Ribo-Seq is especially challenging in bacteria because, unlike in yeast and eukaryotes, bacteria have a broad size distribution of ribosome-protected footprints, ranging from 15-40 nucleotides^3^. The size range that should be selected can vary across Ribo-Seq protocols; at present, most Ribo-Seq experiments on bacteria target a size range of 15 to 45 nucleotides, as was used by Li et al, 2014^1^. Hence, bacterial ribosome profiling protocols must adopt less stringent size selection to comprehensively capture biologically relevant, actively translated RNAs.

Selecting for a wider range of fragments likely enables RNA contaminants that are not associated with the ribosome to persist in bacterial Ribo-Seq libraries, including structured non-coding RNAs (ncRNAs)^4^. These ‘contaminant’ reads have not been thoroughly investigated and are regularly discarded in computational analyses^4,5^. However, curiously, these contaminant ncRNAs have passed the size exclusion steps and survived the MNase treatment, likely without protection from the ribosome in order to be represented in the Ribo-Seq library. We hypothesized that some of these contaminants survive MNase treatment because they are protected from lysis by virtue of their secondary structures. This hypothesis is conceptually similar to one utilized in the method FragSeq^6^; however, FragSeq utilizes a different enzyme, nuclease P1, for fragmentation and aims to probe specific secondary structures of RNA via fragmentation patterns *in vitro^6^*. Here, we propose that instead of disregarding these contaminant signals in Ribo-Seq libraries, the MNase treatment, much like nuclease P1 in FragSeq^6^, may provide valuable insight in identifying RNA structures *in vivo*.

To test if fragmentation correlates with the structural accessibility of RNAs, we visualized the fragmentation pattern across a highly transcribed structured RNA, *ssrS,* native to *E. coli* (Figure 1). The structure of *ssrS* in *E. coli* has been previously validated^8–10^. Focusing only on the 5’ and 3’ ends of reads, representing where MNase fragmentation of the RNA occured, we find that ends of Ribo-Seq reads were overrepresented specifically at junctions between structured and unstructured regions of *ssrS*. This association was reproducibly observed across studies - in our Ribo-Seq experiments on *E. coli* MG1655 (Figure 1A), similar experiments performed by Li et al in 2014^1^, and from MetaRibo-Seq experiments carried out on a fecal sample containing a clinical *E. coli* strain, referred to in a previous manuscript as Sample E^11^. Importantly, this fragmentation pattern was not reproduced in RNA-seq libraries that were not exposed to MNase digestion^1^ (Figure 1D). Therefore, it is likely that *in vivo* secondary structures within *ssrS* protect it from MNase digestion in Ribo-Seq protocols.

**Figure 1:**
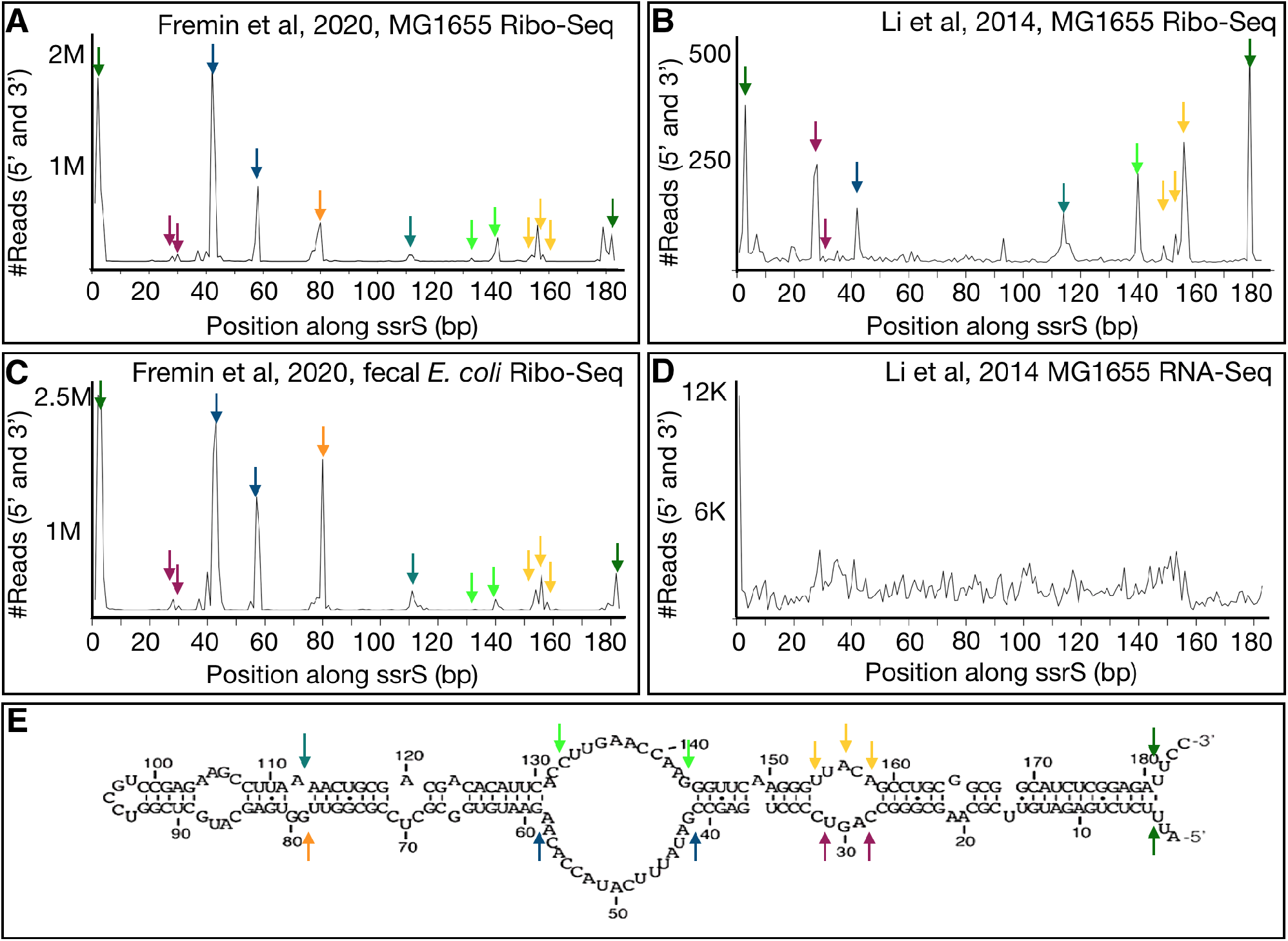
Ribo-Seq fragmentation patterns of ssrS suggest RNA secondary structures protect it from MNase. (A) Quantification of the 3’ and 5’ ends of Fremin et al, 2020 Ribo-Seq reads mapping to *ssrS* in *E. coli* MG1655. Arrows indicate peaks in signal. (B) Quantification of the 3’ and 5’ ends of Li et al, 2014 Ribo-Seq reads mapping to *ssrS* in *E. coli* MG1655. (C) Quantification of the 3’ and 5’ ends of Fremin et al, 2020 MetaRibo-Seq reads mapping to *ssrS* in *E. coli* within a fecal sample. (D) Quantification of the 3’ and 5’ ends of Li et al, 2014 RNA-Seq reads mapping to *ssrS* in *E. coli* MG1655. (E) Characterized structure of *ssrS* in *E. coli*. This structure diagram was taken from previous work^8–10^. Arrows indicate relative positions comparing line graphs (A-D) to this structure diagram.

To further test the broad applicability of data from Ribo-Seq experiments to elucidate the *in vivo* structure of ncRNAs across bacterial species, we next turned our attention to CRISPR arrays from *Ruminococcus.* We hypothesized that since direct repeats are the only structured regions of RNA in CRISPR arrays, only these would survive MNase treatment and therefore be represented in Ribo-Seq data. To test this, we inspected MetaRibo-Seq signal distribution along CRISPR arrays, and found a strong enrichment for structured repeats in the CRISPR arrays (Figure 2). For example, a CRISPR array containing 18 repeats in *Ruminococcus lactaris*, a human gut commensal, contained Ribo-Seq signal specific to each of the 18 repeats in the array (Figure 2B). This suggested that MNase was able to digest spacer regions in these CRISPR arrays but was unable to digest the structured direct repeat regions. Notably, this reinforces our hypothesis that structured regions of ncRNAs escape MNase digestion and therefore are represented in Ribo-Seq experiments.

**Figure 2:**
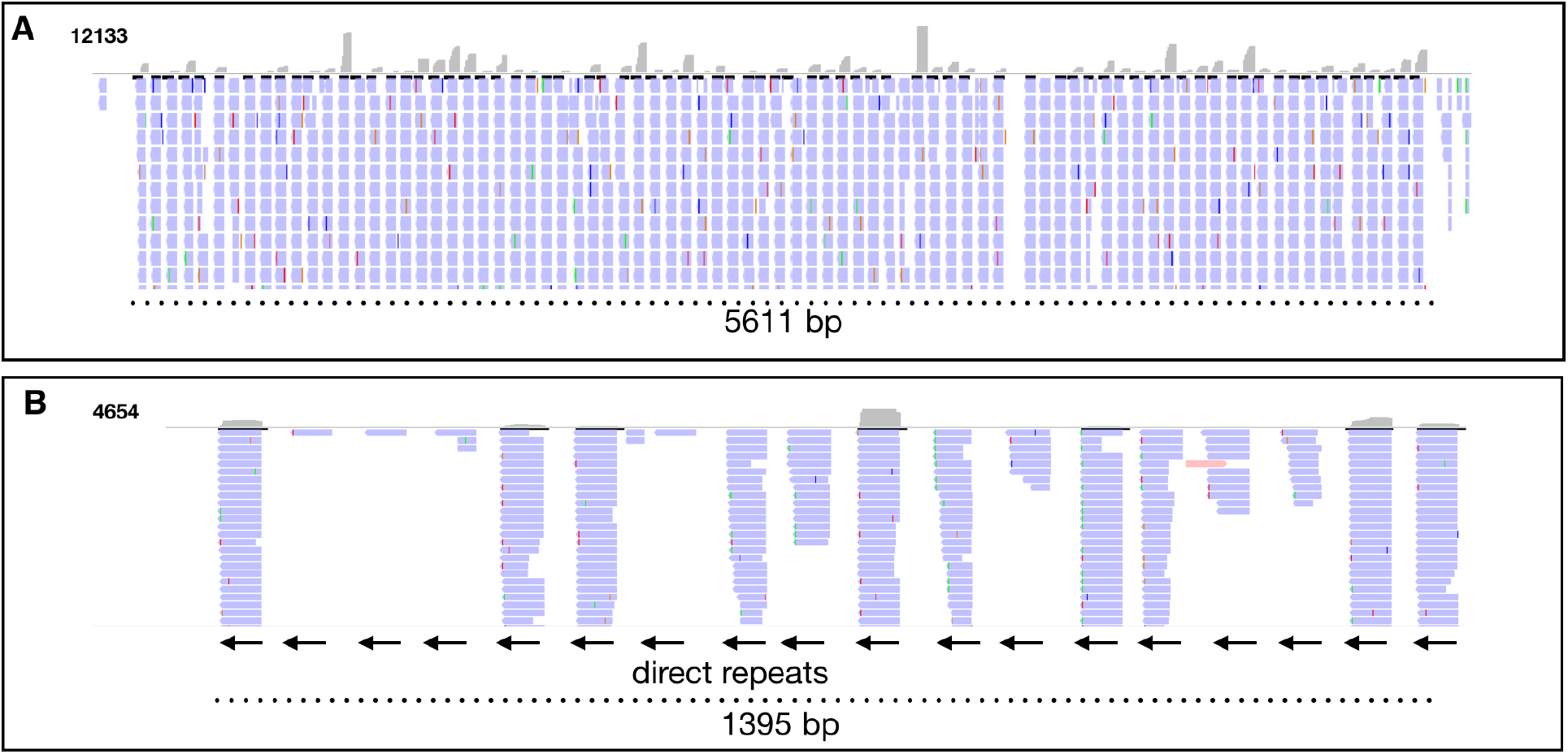
MetaRibo-Seq signal across CRISPR arrays in two gut commensals suggests secondary structures of direct repeats protect it from MNase. (A) Ribo-Seq signal across a CRISPR array containing 84 repeats, predicted by minCED^12^. This is found in *Ruminococcus sp. UNK.MGS-30*. For reference, this was predicted from Sample C in previous work^11^. (B) Ribo-Seq signal across an 18 repeat CRISPR array in *Ruminococcus lactaris*, also predicted by minCED^12^. For reference, this was predicted from Sample A in previous work^11^. Arrows indicate direct repeats.

Importantly, we find that observing Ribo-Seq signals across a structured RNA is not a rare phenomenon limited to these two examples. In fact, the majority of ncRNAs in *E. coli* produce a Ribo-Seq signal (Table S1). We quantified Ribo-Seq and RNA-Seq reads across 65 known ncRNAs in *E. coli* MG1655. All of these ncRNAs were found to be transcribed (RPKM > 10) in RNA-Seq data from Li et al, 2014^1,2^. 61 of the 65 known ncRNAs (94%) produced a Ribo-Seq signal (RPKM > 10) in Ribo-Seq experiments from Li et al, 2014^1,2^ and in Ribo-Seq of MG1655 *E. coli* performed in our laboratory and recently reported^*11*^ (Table S1). Taken together, this highlights that ‘contaminant’ signals in Ribo-Seq experiments are widespread across different structured RNAs.

While this approach represents and exciting new “repurposing” of exisiting Ribo-Seq data, there are several limitations to using contaminant Ribo-Seq signals to gain insights into the structure of RNAs. First, this method is not designed to study structured RNAs and in fact contains steps to actively filter out such contaminants. Ribo-Seq protocols enrich for ribosomes and restrict RNA sequences to a specific size range - therefore, many fragments of RNA that are of structural interest are experimentally removed. Further, this process of eliminating RNA fragments results in a fragmentation profile that is incomplete. The absence of a peak in a Ribo-Seq fragmentation profile for a given structured RNA does not imply the specific structure is not there. We refrain from drawing conclusions from the intensity of any given peak as this could be influenced by transcript abundance, MNase specificity, fragment length, and strength of RNA structure during MNase treatment. Methods like FragSeq^6^ and Shape-Seq^13^ will undoubtedly be more sensitive and provide a more comprehensive catalog of structured RNAs. Additionally, MNase may not be the best enzyme for such fragmentation. From a methodological standpoint, Ribo-Seq cannot match the resolution or completeness of existing technologies to probe for the structures of RNAs. As Ribo-Seq protocols continue to improve, the existence of these contaminants will also diminish.

However, there are also several notable strengths to using contaminant Ribo-Seq signals to gain insight into structured RNAs. Currently there is a plethora of Ribo-Seq data, especially with the development of MetaRibo-Seq and ability to capture the ribosome profile of thousands of taxa at once. To our knowledge, no one one has performed a method like FragSeq^6^ or Shape-Seq^13^ on a complex fecal community. Ribo-Seq has the advantage of capturing *in vivo* RNA structures, in high-throughput, and can immediately be applied to the vast existing datasets. Additionally, Ribo-Seq data can be leveraged to identify novel structured RNAs, many of which are yet to be discovered^14^.

In summary, here we demonstrate an approach to repurpose Ribo-Seq reads to learn about secondary structures in RNAs. First, we analyzed the fragmentation pattern of a well established structured RNA, *ssrS*, in *E. coli*. We observed that the ends of Ribo-Seq reads accumulated at junctions between structured and unstructured regions of the *ssrS* RNA, suggesting that the RNA structure is protected against MNase digestion, akin to FragSeq^6^. Second, we inspected the signal distribution along CRISPR arrays in *Ruminococcus lactaris*. We observed that structured repeats within CRISPR arrays^7^ retained Ribo-Seq reads while spacer regions did not retain reads, suggesting that the structure of the direct repeats were protected from MNase. By focusing on these ‘contaminants’ in Ribo-Seq data, we specifically addressed why they exist in this data type and how they can be useful to researchers interested in the *in vivo* structure of RNAs.

## Acknowledgements

We would like to thank Aravind Natarajan for feedback on the manuscript. Sequencing costs were supported via NIH S10 Shared Instrumentation Grant (1S10OD02014101) and Damon Runyon Clinical Investigator Award to ASB, Stanford ADRC grant # P50AG047366. B.J.F is supported by a National Science Foundation Graduate Research Fellowship DGE-114747 and the Stanford Center Computation, Evolutionary, and Human Genomics fellowship.

## Materials and Methods

### Data Download

Reads from all samples used are publicly available. The in-house generated data can be found under this accession: PRJNA510123^11,14^. Ribo-Seq and RNA-Seq for *E. coli* generated by Li et al, 2014 can be found under this bioproject: PRJNA232843^1^.

### Genome Annotation

CRISPR arrays were predicted from reference genomes using minCED^4^ as a part of Prokka v1.12^15^.

### Read Mapping

Reads were trimmed with trim galore version 0.4.0 using cutadapt 1.8.1^16^ with flags –q 30 and – illumina. Reads were mapped to the annotated assemblies using bowtie version 1.1.1^17^. Reads were counted using bedtools^18^ multicov. The 5’ and 3’ positions of reads were determined using bedtools^18^ genomecov. IGV^19^ was used to visualize coverage. Reads per Kilobase Million (RPKM) calculations were performed using in house scripts.

## References

1. Li, G.-W., Burkhardt, D., Gross, C. & Weissman, J. S. Quantifying absolute protein synthesis rates reveals principles underlying allocation of cellular resources. Cell 157, 624–635 (2014).

2. Ingolia, N. T., Ghaemmaghami, S., Newman, J. R. S. & Weissman, J. S. Genome-wide analysis in vivo of translation with nucleotide resolution using ribosome profiling. Science 324, 218–223 (2009).

3. Mohammad, F., Green, R. & Buskirk, A. R. A systematically-revised ribosome profiling method for bacteria reveals pauses at single-codon resolution. Elife 8, (2019).

4. Guttman, M., Russell, P., Ingolia, N. T., Weissman, J. S. & Lander, E. S. Ribosome Profiling Provides Evidence that Large Noncoding RNAs Do Not Encode Proteins. Cell vol. 154 240–251 (2013).

5. Brar, G. A. & Weissman, J. S. Ribosome profiling reveals the what, when, where and how of protein synthesis. Nat. Rev. Mol. Cell Biol. 16, 651–664 (2015).

6. Underwood, J. G. et al. FragSeq: transcriptome-wide RNA structure probing using high-throughput sequencing. Nat. Methods 7, 995–1001 (2010).

7. Kunin, V., Sorek, R. & Hugenholtz, P. Evolutionary conservation of sequence and secondary structures in CRISPR repeats. Genome Biol. 8, R61 (2007).

8. Reiche, K. & Stadler, P. F. RNAstrand: reading direction of structured RNAs in multiple sequence alignments. Algorithms Mol. Biol. 2, 6 (2007).

9. Panchapakesan, S. S. S. & Unrau, P. J. E. coli 6S RNA release from RNA polymerase requires σ70 ejection by scrunching and is orchestrated by a conserved RNA hairpin. RNA 18, 2251–2259 (2012).

10. Cavanagh, A. T., Sperger, J. M. & Wassarman, K. M. Regulation of 6S RNA by pRNA synthesis is required for efficient recovery from stationary phase in E. coli and B. subtilis. Nucleic Acids Research vol. 40 2234–2246 (2012).

11. Fremin, B. J. & Bhatt, A. S. Metagenome-wide measurement of protein synthesis in the human fecal microbiota using MetaRibo-Seq. doi:10.1101/482430.

12. Bland, C. et al. CRISPR recognition tool (CRT): a tool for automatic detection of clustered regularly interspaced palindromic repeats. BMC Bioinformatics 8, 209 (2007).

13. Watters, K. E., Abbott, T. R. & Lucks, J. B. Simultaneous characterization of cellular RNA structure and function with in-cell SHAPE-Seq. Nucleic Acids Res. 44, e12 (2016).

14. Fremin, B. J. & Bhatt, A. S. A combined RNA-Seq and comparative genomics approach identifies 1,085 candidate structured RNAs expressed in human microbiomes. doi:10.1101/2020.03.31.018887.

15. Seemann, T. Prokka: rapid prokaryotic genome annotation. Bioinformatics 30, 2068–2069 (2014).

16. Martin, M. Cutadapt removes adapter sequences from high-throughput sequencing reads. EMBnet.journal 17, 10 (2011).

17. Langmead, B., Trapnell, C., Pop, M. & Salzberg, S. L. Ultrafast and memory-efficient alignment of short DNA sequences to the human genome. Genome Biol. 10, R25 (2009).

18. Quinlan, A. R. BEDTools: The Swiss-Army Tool for Genome Feature Analysis. Curr. Protoc. Bioinformatics 47, 11.12.1–34 (2014).

19. Thorvaldsdottir, H., Robinson, J. T. & Mesirov, J. P. Integrative Genomics Viewer (IGV): high-performance genomics data visualization and exploration. Brief. Bioinform. 14, 178–192 (2012).

